## supplementary code and tables for "Dimorphic Neural Network Architecture Prioritizes Sexual-related Behaviors in Male C.elegans": Supplementary file 1.py

```
import copy

# 线虫及突触
nematode_type = 'hermaphrodite'  # 'hermaphrodite' 或 'male'
synapse_type = 'all'  # 'chemical' 或 'gap_symmetric' 或 'all'

if nematode_type == 'hermaphrodite':
    neuron_list = []
    with open('./connection_data/hermaphrodite_neuron_list.csv') as file_object:
        for line in file_object:
            neuron_list.append(line.replace('\n', ''))
    neuron_number = len(neuron_list)
    pharynx_neuron = neuron_list[0:14]
    sensory_neuron = neuron_list[14:53]
    inter_neuron = neuron_list[53:98]
    motor_neuron = neuron_list[98:122]
    neuron_connection = [[0 for _ in range(neuron_number)] for _ in range(neuron_number)]
    if synapse_type == 'chemical':
        with open('./connection_data/hermaphrodite_chemical_connection.csv') as file_object:
            i = 0
            for line in file_object:
                a = line.split(',')
                for j in range(neuron_number):
                    neuron_connection[i][j] = int(a[j].replace('\n', ''))
                i += 1
    elif synapse_type == 'gap_symmetric':
        with open('./connection_data/hermaphrodite_gap_symmetric_connection.csv') as file_object:
            i = 0
            for line in file_object:
                a = line.split(',')
                for j in range(neuron_number):
                    neuron_connection[i][j] = int(a[j].replace('\n', ''))
                i += 1
    elif synapse_type == 'all':
        with open('./connection_data/hermaphrodite_all_connection.csv') as file_object:
            i = 0
            for line in file_object:
                a = line.split(',')
                for j in range(neuron_number):
                    neuron_connection[i][j] = int(a[j].replace('\n', ''))
                i += 1
    else:
        raise ValueError(
            '未定义的连接类型。允许输入的连接类型为chemical或gap_symmetric或all，但实际的输入为{}'.format(synapse_type)
        )
elif nematode_type == 'male':
    neuron_list = []
    with open('./connection_data/male_neuron_list.csv') as file_object:
        for line in file_object:
            neuron_list.append(line.replace('\n', ''))
    neuron_number = len(neuron_list)
    pharynx_neuron = neuron_list[0:14]
    sensory_neuron = neuron_list[14:53] + neuron_list[120:121] + neuron_list[122:149]
    inter_neuron = neuron_list[53:98] + neuron_list[121:122] + neuron_list[149:161]
    motor_neuron = neuron_list[98:120]
    neuron_connection = [[0 for _ in range(neuron_number)] for _ in range(neuron_number)]
    if synapse_type == 'chemical':
        with open('./connection_data/male_chemical_connection.csv') as file_object:
            i = 0
            for line in file_object:
                a = line.split(',')
                for j in range(neuron_number):
                    neuron_connection[i][j] = int(a[j].replace('\n', ''))
                i += 1
    elif synapse_type == 'gap_symmetric':
        with open('./connection_data/male_gap_symmetric_connection.csv') as file_object:
            i = 0
            for line in file_object:
                a = line.split(',')
                for j in range(neuron_number):
                    neuron_connection[i][j] = int(a[j].replace('\n', ''))
                i += 1
    elif synapse_type == 'all':
        with open('./connection_data/male_all_connection.csv') as file_object:
            i = 0
            for line in file_object:
                a = line.split(',')
                for j in range(neuron_number):
                    neuron_connection[i][j] = int(a[j].replace('\n', ''))
                i += 1
    else:
        raise ValueError(
            '未定义的连接类型。允许输入的连接类型为chemical或gap_symmetric或all，但实际的输入为{}'.format(synapse_type)
        )
else:
    raise ValueError(
        '未定义的线虫类型。允许输入的线虫类型为hermaphrodite或male，但实际的输入为{}'.format(nematode_type)
    )

# 神经元参数
c = 1.0  # 细胞膜电容
g_l = 0.03  # 泄漏电导
e_l = -54.387  # 泄漏反转电位
g_na = 120.  # 钠离子通道电导
e_na = 50.  # 钠离子通道反转电位
g_k = 36.  # 钾离子通道电导
e_k = -77.  # 钾离子通道反转电位

# 神经元膜电位
syn_rate = 100.

# 模拟时间及步长
total_time = 10
each_step = 0.01
total_step = int(total_time / each_step)

# 输入电压
volt_set = 60.
for neuron in range(neuron_number):
    neuron_potential = [-60. for _ in range(neuron_number)]
    neuron_potential[neuron] = volt_set

    # 运行
    with open('./d/{}_{}_output.csv'.format(neuron_list[neuron], nematode_type), 'a') as file_object:

        for i in range(total_step):

            # 导出数据
            for j in range(neuron_number):
                if j == neuron_number - 1:
                    file_object.write(str(neuron_potential[j]) + '\n')
                else:
                    file_object.write(str(neuron_potential[j]) + ',')

            neuron_potential_cash = copy.deepcopy(neuron_potential)

            for j in range(neuron_number):

                if j == neuron:
                    neuron_potential_cash[j] = volt_set
                    continue

                sy_current = 0
                for k in range(neuron_number):
                    if neuron_connection[k][j] != 0:
                        sy_current = sy_current + neuron_connection[k][j] * syn_rate * (
                                    neuron_potential[k] - neuron_potential[j])
                k1 = (-1 * g_l * (neuron_potential[j] - e_l) - g_k * (neuron_potential[j] - e_k) - g_na * (
                        neuron_potential[j] - e_na) + sy_current) / c

                neuron_potential[j] = neuron_potential_cash[j] + k1 * each_step * 0.5

                sy_current = 0
                for k in range(neuron_number):
                    if neuron_connection[k][j] != 0:
                        sy_current = sy_current + neuron_connection[k][j] * syn_rate * (
                                    neuron_potential[k] - neuron_potential[j])
                k2 = (-1 * g_l * (neuron_potential[j] - e_l) - g_k * (neuron_potential[j] - e_k) - g_na * (
                        neuron_potential[j] - e_na) + sy_current) / c

                neuron_potential[j] = neuron_potential_cash[j] + k2 * each_step * 0.5

                sy_current = 0
                for k in range(neuron_number):
                    if neuron_connection[k][j] != 0:
                        sy_current = sy_current + neuron_connection[k][j] * syn_rate * (
                                    neuron_potential[k] - neuron_potential[j])
                k3 = (-1 * g_l * (neuron_potential[j] - e_l) - g_k * (neuron_potential[j] - e_k) - g_na * (
                        neuron_potential[j] - e_na) + sy_current) / c

                neuron_potential[j] = neuron_potential_cash[j] + k3 * each_step

                sy_current = 0
                for k in range(neuron_number):
                    if neuron_connection[k][j] != 0:
                        sy_current = sy_current + neuron_connection[k][j] * syn_rate * (
                                    neuron_potential[k] - neuron_potential[j])
                k4 = (-1 * g_l * (neuron_potential[j] - e_l) - g_k * (neuron_potential[j] - e_k) - g_na * (
                        neuron_potential[j] - e_na) + sy_current) / c

                neuron_potential[j] = neuron_potential_cash[j]
                neuron_potential_cash[j] = neuron_potential_cash[j] + (k1 + 2 * k2 + 2 * k3 + k4) * each_step / 6.

            neuron_potential = copy.deepcopy(neuron_potential_cash)
            print(neuron_list[neuron] + '完成' + str(round(i / total_step * 100, 2)) + '%')
```
